# Pescoids and chimeras to probe early evo-devo in the fish *Astyanax mexicanus*

**DOI:** 10.1101/2020.10.06.328500

**Authors:** Jorge Torres-Paz, Sylvie Rétaux

## Abstract

The fish species *Astyanax mexicanus* with its sighted and blind eco-morphotypes has become an original model to challenge vertebrate developmental evolution. Recently, we demonstrated that phenotypic evolution can be impacted by early developmental events starting from the production of oocytes in the fish ovaries. *A. mexicanus* offers an amenable model to test the influence of maternal effect on cell fate decisions during early development, yet the mechanisms by which the information contained in the eggs is translated into specific developmental programs remain obscure due to the lack of specific tools in this emergent model. Here we describe methods for the generation of gastruloids from yolkless-blastoderm explants to test the influence of embryonic and extraembryonic properties on cell fate decisions, as well as the production of chimeric embryos obtained by intermorph cell transplantations to probe cell autonomous or non-autonomous processes. We show that *Astyanax* gastruloids have the potential to recapitulate the main ontogenetic events observed in intact embryos, including the internalization of mesodermal progenitors and eye development, as followed with *Zic:GFP* reporter lines. In addition, intermorph cell grafts resulted in proper integration of exogenous cells into the embryonic tissues, with lineages becoming more restricted from mid-blastula to gastrula. The implementation of these approaches in *A. mexicanus* will bring new light on the cascades of events, from the maternal pre-patterning of the early embryo to the evolution of brain regionalization.

## Introduction

Emergent model organisms offer novel possibilities to unravel specific questions related to developmental and cell biology. However, some limitations, inherent to each animal system, often render difficult the implementation of novel methodologies.

*Astyanax mexicanus* species has thrived as an emergent model organism for evolutionary developmental biology studies (Jeffery, 2020; Torres-Paz et al., 2018). Its success in this field is due to the existence of two markedly different eco-morphotypes within the same species. *A. mexicanus* comprises river-dwelling fish populations, “surface fish”, and several populations adapted to the life in caves in complete and permanent darkness, “cavefish”. During cave adaptation, striking morpho-functional modifications occurred. Compared to the surface fish, the cave-adapted morphs have completely lost their eyes and pigmentation. In addition, some constructive traits have also emerged such as larger olfactory organs and more numerous facial neuromasts and taste buds, which probably contribute to a sensory compensation for the loss of the visual system (Bibliowicz et al., 2013; Blin et al., 2018; Hinaux et al., 2016; Varatharasan et al., 2009; Yoshizawa et al., 2010). Most of the morphological differences observed in the nervous system of *A. mexicanus* morphotypes have an early embryonic origin (Hinaux et al., 2016; Pottin et al., 2011; Rétaux et al., 2016; Yamamoto & Jeffery, 2000). In fact, recent evidence has shown that maternal determinants, present in the oocyte before fertilization and before zygotic developmental programs are initiated, have an important contribution to later phenotypes (Ma et al., 2020, Ma et al., 2018; Torres-Paz et al., 2019). Indeed, any differential composition of maternal determinants in the eggs is susceptible to lead to changes in early developmental events, such as activation of the zygotic genome, embryonic patterning and establishment of signaling centers, thus affecting later ontogenetic processes.

In fish, the extraembryonic yolk cell (of maternal origin) is an important source of inductive signals that pattern the overlying blastoderm, the embryo proper. Asymmetric segregation of maternal determinants leads to the induction of the embryonic organizer in the prospective dorsal side of the blastoderm. This symmetry breaking event will lead to localized production of different morphogens, creating gradients of signaling activity within the developing embryo. The integration of these signals by embryonic cells provides them with positional information and instructs them to follow a particular developmental program (Schier & Talbot, 2005). Thus, changes in the information contained in the oocytes, represented by maternally inherited RNAs and proteins, will affect the subsequent sequence of developmental events. *A. mexicanus* offers a unique model to test the maternal influence on precocious and later development. However, methods to assess the effect of signaling centers (embryonic and extraembryonic) and the potential of cells to respond to these signals have not been developed yet in this model.

Here, we describe the implementation in *A. mexicanus* of methods used in well-established fish models to probe mechanisms of cell/tissue specification during early embryogenesis. First, we have adapted a recent method of embryonic explant culture developed in zebrafish and known as “pescoids” to grow the blastoderm after removal of the extraembryonic yolk cell (Fulton et al., 2020; Schauer et al., 2020). Under these conditions of altered embryonic geometry and physical constraints, the explants are able to recapitulate the main processes observed in intact embryos such as symmetry breaking, germ layer specification and elongation. In addition to previous reports on fish gastruloids, here we found clear indications of mesoderm internalization and eye development. These gastruloids will allow comparative analyses of gene expression in *Astyanax* morphs in the absence of yolk-derived signaling. Second, we have set up the conditions to efficiently achieve inter-morph cell transplantations at matching stages during early embryogenesis. Cell grafting have been widely performed in zebrafish embryos to test cell autonomy and potential during development, as well as to dissect lineage and timing aspects during cell specification. Grafts have also been performed between distinct species such as zebrafish and medaka to study developmental heterochronies (Fuhrmann et al., 2020). In *A. mexicanus*, inter-morphs cell transplantation will allow asking similar questions in a micro-evolutionary context. Further, the implementation of these methodologies to generate gastruloids and chimeric embryos in *A. mexicanus* will help to explore the effect of embryonic and extraembryonic signals in cell decisions during early development.

## Materials and methods

### Fish and embryo collection

Our *A. mexicanus* colonies were obtained in 2004 from the Jeffery laboratory at the University of Maryland, College Park, United States. The surface fish stock derives from rivers in Texas, United States and the cavefish from the Pachón cave in San Luis Potosi, Mexico. Fish were since then maintained on a 12:12 hr light:dark cycle at a temperature of 22°C for cavefish and 26°C for surface fish. Reproductions were induced every other week by changes in water temperature: for cavefish temperature was increased to 26°C, and for surface fish temperature was decreased to 22°C during 3 days followed by an increase to 26°C (Elipot et al., 2014). Fish from both morphotypes spawn regularly the first and second days following the increase in temperature. Here, embryos were obtained exclusively by *in vitro* fertilization in order to ascertain synchronous early development. Embryo dechorionation was performed by enzymatic treatment with Pronase 1mg/mL (Sigma) and embryos were maintained in Embryo Medium (EM) at 24°C. Surface and Pachón cavefish *zic1:GFP* transgenic lines used here were generated previously in the lab (Devos et al., 2019). Animals were treated according to the French and European regulations for handling of animals in research. SR’s authorization for use of *Astyanax mexicanus* in research is 91-116. This work did not necessitate a protocol authorization number from the Paris Centre-Sud Ethic Committee. The animal facility of the Institute received authorization 91272105 from the Veterinary Services of Essonne, France, in 2015.

### Generation of chimeric embryos by cell transplantations

Donor embryos were injected at the 1-cell stage with 3-5nL of 1% Dextran-FITC 10,000 MW (Molecular Probes) and 0.05% Phenol Red (to see the solution) using a FemtoJet (Eppendorf). Glass pipettes for microinjection and cell transplantation were prepared on a Narishige PN-30 puller using borosilicate glass capillary (GC100F15 Harvard Apparatus LTD and B120-69-10 WPI, respectively). Microinjection pipettes were sealed at the tip and broken for opening at the moment of the injection using forceps. Cell transplantations pipettes were prepared in advance, the tip was broken at the desired internal diameter (15-30 µm) and polished using a Micropipettte grinder (Narishige EG-44) at an angle of 35° in order to create a smooth needle-shaped tip. Our cell transplantation system consisted of a holder for the glass pipette (WPI) connected to a 1mL syringe by a Teflon tubing (Narishige). Under a fluorescent macroscope labelled donor cells were aspirated into the tip of the glass pipette filled with EM, and 3-12 cells expelled into the host embryo with gentle pressure. Host embryos were let to develop in EM until fixation. In this study, isotopic and isochronic intermorphs grafts were performed (Surface animal pole cells to Cave animal pole either at blastula or gastrula stages).

### Generation of gastruloids

*A. mexicanus* gastruloids were produced following a recent description in zebrafish (Fulton et al., 2020; **Figure 1A**). Briefly, at the 512-1K cell stage the yolk was carefully removed from embryos using eyebrows knives (**Supplemental video 1**). Blastoderm explants were cultured until the corresponding 10 hours post-fertilization (hpf) at 24°C in L15 medium (Gibco) supplemented with 3% Fetal Bovine Serum (FBS, Biosera). For cultures maintained for longer than 7hrs Penicillin-Streptomycin were added to the culture media (1x dilution, P4333, Sigma).

**Figure 1.**
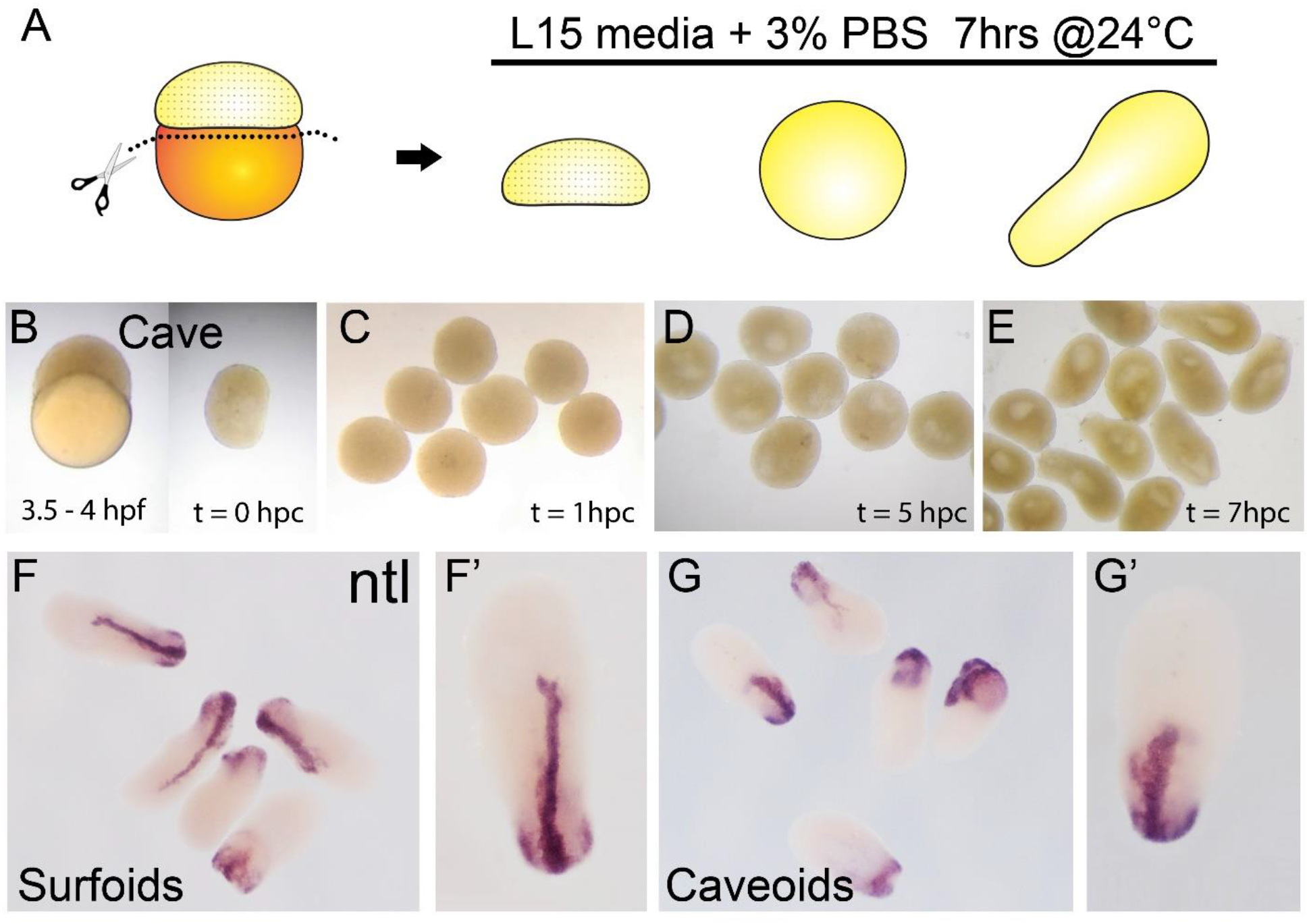
Generation of pescoids in *A. mexicanus*. Procedure for the generation of *Astyanax* pescoids (**A**). Intact cavefish embryo at 1K-cell stage (**B**, left) and after yolk removal (**B**, right). Development of caveoids after 1, 5 and 7 hpc (**C, D** and **E**, respectively). Expression of *ntl* in surfoids (**F, F’**) and caveoids (**G, G’**) after 7 hpc.

### Histology and imaging

Colorimetric ISH was performed as previously described (Pottin et al., 2011). Digoxigenin-labeled riboprobe was prepared using PCR products as templates. *no-tail* (*ntl*) cDNA was obtained from our ESTs library (accession number ARA0ABA99YL22). Procedure for revelation of FITC-labeled donor cells combined with fluorescent *ISH* was adapted from our *ISH* protocol (Alié et al., 2018). Following the hybridization with the Digoxigenin-labeled ISH probe, embryos were first processed for the revelation of the Dextran-FITC grafted cells: they were incubated for 1 hour in blocking solution (Tris 0.1M pH 7.5, NaCl 150mM, Tween-20 0.1% and 5% blocking reagent Roche) and then with POD-conjugated anti-FITC antibody (11426346910; Roche, 1/400) diluted in blocking solution overnight. Embryos were washed in PBS/Tween 0.1% (PBST) 10 times for 10 minutes each time and incubated for 30 min at room temperature with TAMRA-tyramide 1/1000. Peroxidase activity was activated by H2O2 (0.003%, Sigma) for 1 hr and samples were washed again 10 times for 10 minutes in PBST. Revelation of the dig-labeled ISH probe was then performed using an anti-Digoxigenin antibody coupled to POD (11207733910; Roche, 1/400) and revealed using a FITC-tyramide (1/400). After several washes in PBST embryos were stained with DAPI (10236276001, Sigma) at a final concentration of 1 mg/ml, overnight at 4°C, and washed in PBS before dissection and mounting (Vectashield, Vector Laboratories).

Immunohistochemistry was performed as previously described (Blin et al., 2018) using a primary anti-GFP antibody at a dilution 1/500 (GFP-1020, Aves Labs) and a secondary Alexa Fluor 488-coupled antibody at a dilution 1/500 (Anti-chicken, A-11039, Invitrogen). Fluorescently labeled embryos were counterstained with DAPI. Embryos stained by colorimetric ISH were imaged on a Nikon AZ100 multizoom macroscope. Confocal acquisitions were done on a Leica-SP8 confocal microscope using the Leica Application Suite software. Images were processed on ImageJ.

## Results and discussion

### Generation of Astyanax gastruloids

Recent advances in the field of gastruloids have highlighted the robustness of animal development and the key steps taking place during this process. In zebrafish pescoids (yolk-less blastoderm explants) the main aspects of early development observed in intact embryos are recapitulated. Symmetry breaking, axis elongation and neural specification occur despite the absence of extraembryonic signaling (Fulton et al., 2020; Schauer et al., 2020). In fish, these events are controlled maternally (Marlow, 2020; Solnica-Krezel, 2020) and *A. mexicanus* with its two morphotypes has become an excellent model to challenge the role of maternal determinants in embryogenesis (Ma et al., 2020, 2018; Torres-Paz et al., 2019).

Pescoids derived from embryos of both *Astyanax* surface and cavefish embryos (surfoids and caveoids, respectively) developed similarly to those described in zebrafish. After removal of the vitellus at the 256-2K cell stage (**Figure 1B**), blastoderm explants sealed the wound and became rounded during the first hour post culture (hpc, **Figure 1C**). Then, during the next 3-4 hours in culture a cavity was formed (**Figure 1C**), which may correspond to a “blastocoel” (Schauer et al., 2020). After 5 hpc, the first signs of axis elongation were observed, similarly in terms of timing in surfoids and caveoids (not shown). Cultures were stopped after 7 hpc, the time corresponding to the tailbud stage in intact embryos, and gastruloids were fixed for further processing and imaging. The extent of elongation at the end of the culture was variable between pescoids and fluctuated similarly in the two morphs: gastruloid elongation was observed in 79% of surfoids and 84% of caveoids. The shape of the elongated gastruloids was asymmetrical and pear-shaped, with a narrow tip at one end and a larger rounded form at the opposite extremity. We studied the expression of the mesodermal marker *ntl* in *Astyanax* pescoids and found a pattern strikingly reminiscent of the developing notochord observed in intact fish embryos (**Figure 1F, G**). Microscopic observations suggested that cells expressing *ntl* had been internalized, an aspect of fish gastruloids that may have been overlooked. We compared the expression of *ntl* in confocal acquisitions after fluorescent ISH in intact embryos at tailbud stage and in pescoids at equivalent stages (**Figure 2**). In confocal reconstructed sections of pescoids and control embryos, we observed similar organization of *ntl* expressing cells, always underneath an overlying layer of superficial cells (**Figure 2**). Thus, these data confirm a conserved internalization process of axial mesoderm in explants despite the absence of vitellus. These observations highlight the robustness of cellular processes during vertebrate gastrulation.

**Figure 2.**
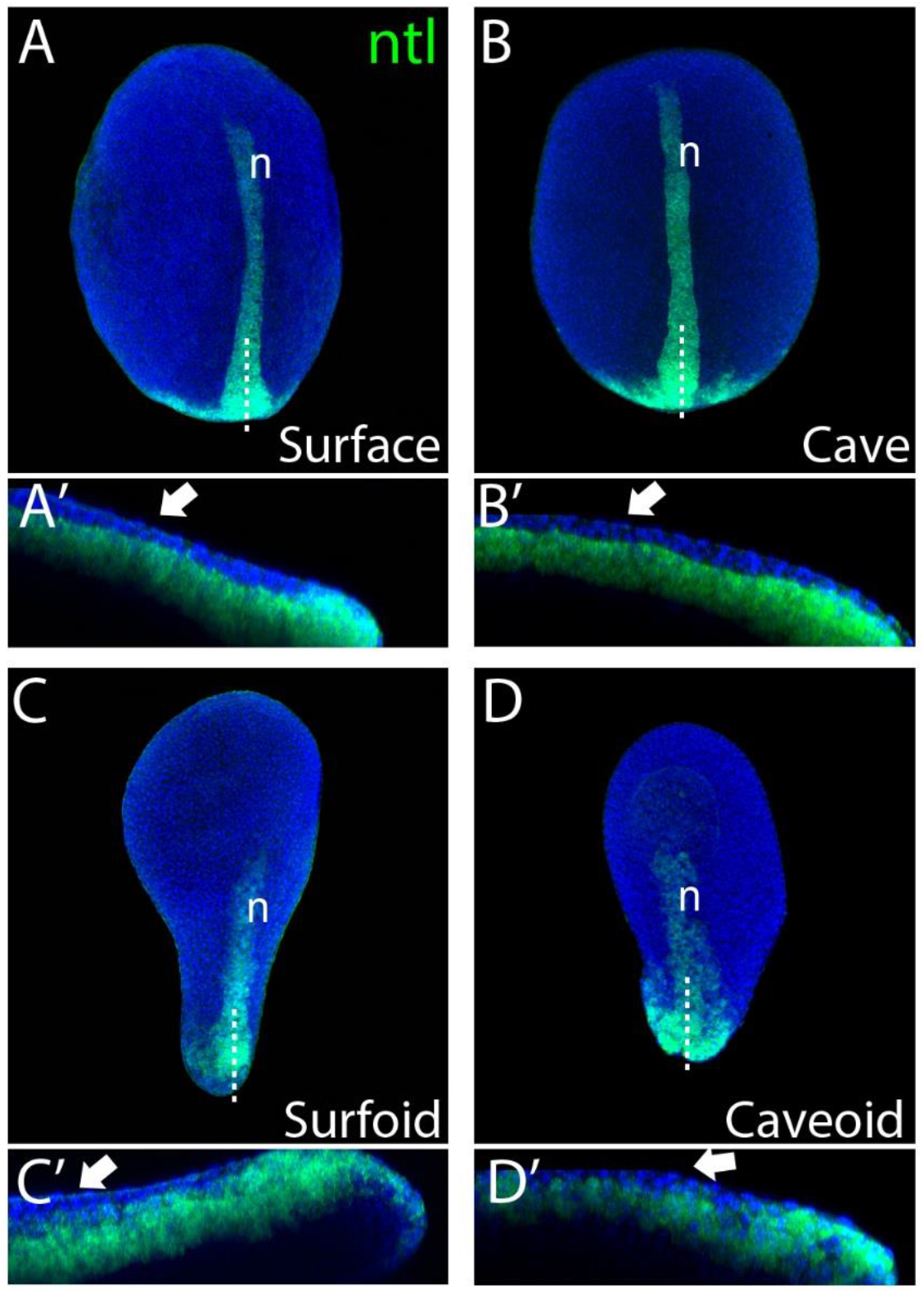
Internalization of mesoderm in *Astyanax* pescoids. Expression of *ntl* in intact surface and cavefish embryos at 10 hpf (**A** and **B**, respectively), and in surfoid and caveoid after 7 hpc (**C** and **D**, respectively). Embryos are oriented in dorsal views, anterior to the top. Bottom images are confocal reconstructions at the indicated level (dotted lines), anterior to the left and dorsal to the top. The white arrows indicate ectodermal cells overlying ntl –expressing mesodermal cells.

Studies in zebrafish have shown that asymmetric translocation of maternal determinants leading to the dorsal determination occur as early as the 16-32 cell stage (Jesuthasan & Strähle, 1997). The activation of the zygotic genome, a process dependent on the maternal transcriptomic machinery, starts around the 64 cell stage (Chan et al., 2019), whereas clear zygotic transcription is observed from the 512 cell stage. Concurrently, at this stage, called the mid-blastula transition, embryonic cell cycles become asynchronous and the extra-embryonic yolk syncytial layer (YSL) is formed at the interphase between the yolk and the blastoderm (Kimmel et al., 1995). Similarly to zebrafish, in *Astyanax* the mid-blastula transition takes place around the 512-1K cell stage (Hinaux et al., 2011). Here, the explant experiments were performed between the 256-2K cell stage, i.e., the end of the maternal-to-zygotic transition, thus it must be taken into account that maternally-derived pre-patterning already exists in the cultured blastoderms. Consequently, in order gain insights into the temporal sequence of maternal patterning events, the generation of earlier explants must be considered. The comparative analysis of earlier explants in the two *Astyanax* morphs will allow dissecting precisely the timing and the impact of the maternal contributions to developmental evolution. Recently, it was shown in zebrafish that the extraembryonic YSL layer does not form properly in yolkless-explants and that correctly developing pescoids can be obtained even from very precocious 64 cell stage embryos (Schauer et al., 2020). This indicates that the gastruloids are able to develop into embryo-like structures in the absence YSL-derived signals (Chen & Kimelman, 2000; Rodaway et al., 1999). Conversely, animal caps explants are able to develop into structures similar to pescoids only if Nodal and downstream planar cell polarity signaling pathways are active (Williams & Solnica-Krezel, 2020), indicating that Nodal activity in pescoids must come from marginal cells. Nodal signaling is necessary for the induction of endomesodermal fates at the blastoderm margin (Schier et al., 1997; Vopalensky et al., 2018). Visual inspection of the mesodermal marker *ntl* expression in our pescoids at the different states of elongation clearly shows that the point where *ntl*-expressing cells are internalized corresponded to the marginal zone, where the wound closed (not shown). Thus, the wounded margin in the blastoderm explants would be topologically and functionally equivalent to the blastopore in intact embryos, being both the source of Nodal signaling and the point where endomesoderm is internalized. Given the previously described differences in organizer formation and mesoderm internalization between the two *Astyanax* morphs embryos (Torres-Paz et al 2019), pescoids will prove powerful tools to study the origin and outcomes of these processes.

### Eye development in Astyanax pescoids

Our observations of axial elongation and the overtly normal formation of the notochord in pescoids (**Figure 2**) made us wonder if eye development could occur in these yolkless explants. We recently generated *zic1:GFP* transgenic knock-in reporter lines in both *Astyanax* morphotypes in order to visualize eye morphogenesis (Devos et al., 2019). We took advantage of these transgenic fish to evaluate directly eye development in pescoids. Blastoderm explants from *zic1:GFP* embryos were let to develop *in vitro* until the equivalent of 24 hpf, i.e., when the optic cups are formed in surface and cave embryos (**Figure 3A, B**). Despite some body axes malformation in these explants (not shown), we observed GFP reporter expression in discrete regions within the prospective pescoid head in some samples (**Figure 3C-E**). Strikingly, we were able to identify distinct GFP expression domains that we interpreted as corresponding to the optic (arrows, **Figure 3**) and telencephalic (“t”, **Figure 3**) *Zic1*-expressing tissues, respectively. The extension of the optic domain delimited by reporter expression varied importantly, in the two morphotypes. Surprisingly, we found in some of our pescoids some degree of retinal morphogenesis (**Figure 3E, E1-E4**), with clear indications of epithelial folding (arrow, **Figure 3E**) and neural tissue contacting adjacent non-neural ectoderm (arrowhead, **Figure E1-E-4**). Our observations highlight the robustness of developmental processes occuring in gastruloids.

**Figure 4.**
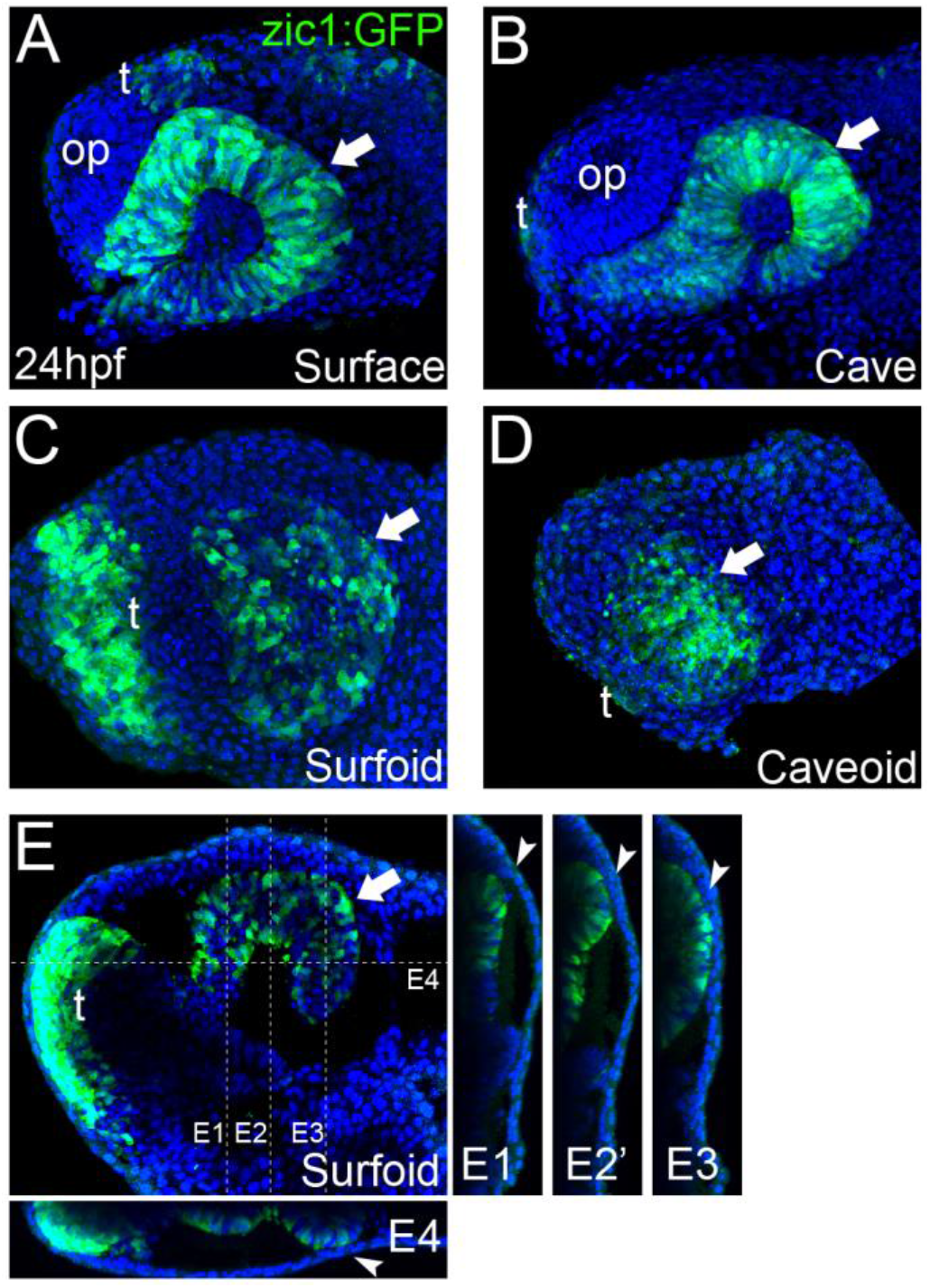
Eye development in *Astyanax* pescoids. GFP-reporter expression in *zic1:GFP* transgenic surface and cavefish embryos (**A** and **B**, respectively), and surfoid (**C** and **E**) and caveoids (**D**) at 24 hpf. Embryos and pescoids are oriented in lateral views, anterior to the left. Arrows indicate the optic tissue; op, olfactory placode; t, telencephalon. Confocal reconstructions at the indicated levels in **E** are shown in **E1, E2, E3** and **E4**. Arrowhead in **E1-E4** indicate the contact zone between neural and non-neural ectoderm.

Eye development starts with the specification eye precursor cells in the anterior neural plate (Cavodeassi et al., 2005; Zuber et al., 2003) followed by complex morphogenesis process that involves coordinated movements within the neural plate (Cavodeassi et al., 2013; Ivanovitch et al., 2013) that are instructed by midline signaling (García-Calero, 2008; Gordon et al., 2018; Macdonald et al., 1995). Our observations of eye-like embryonic tissue in *Astyanax* pescoids highlight the potential of gastruloid systems to engage into complex morphogenesis. The wide spectrum of optic phenotypes observed in our pescoids will allow a better understanding of minimal requirements for particular developmental processes and the interdependency of different embryonic tissues during morphogenesis.

### Generation of chimeric embryos by cell grafting

Cell transplantation methodologies have been widely used in zebrafish to test cell-autonomy processes during embryogenesis in different experimental contexts (Cavodeassi et al., 2013; Fauny et al., 2009; Giger et al., 2016). *A. mexicanus* with its two eco-morphotypes offers a great opportunity to test autonomy during cell fate specification through intermorph transplantations. Hence, elegant transplantation experiments of the lens from one morph into the optic cup of the other morph at larval stages have revealed the autonomy of the cavefish lens apoptotic process and its role in cavefish eye degeneration (Yamamoto & Jeffery, 2000). Neural crest cell transplantations have been performed as well at 2dpf to address their role in cavefish pigmentation defect (Yoshizawa et al., 2018). However, the *Astyanax* model has not succeeded yet for early cell transplantations aimed at studying precocious embryogenesis, mainly due to technical challenges that must be circumvented. A major challenge is the simultaneous collection of embryos of both morphotypes at equivalent developmental stage. *A. mexicanus* reproduce in the dark (Simon et al., 2019), thus in order to obtain early developing embryos to work with during the day, we inverted the circadian cycle of fish in a special fish room dedicated to reproduction in our facility. In addition, to obtain early embryos developing synchronously, *in vitro* fertilizations must be performed using ready-to-spawn females, hence mating behavior were monitored during the days following the induction of reproductions with an infrared camera in the circadian-inverted fish room. Inductions of surface and cavefish were then coordinated in order to find spawning females of the two morphs at the same time.

In this work surface fish embryos were labeled with dextran-FITC at the one-cell stage (donors), and cells were grafted isochronically into unlabeled cavefish embryos (hosts) at two developmental time points, the mid-blastula transition (512-2K cell stage) and the onset of gastrulation (30-50% epiboly) (**Figure 4A**). As a source of donor cells we choose the embryonic animal pole, in order to compare to fate maps studies performed in zebrafish and showing that ectodermal precursors (including neurectoderm) arise from this field, whereas endomesodermal precursors derive from more marginal cells (Schier & Talbot, 2005; Woo & Fraser, 1997). After transplantation of fluorescently labeled cells (**Figure 4B**), embryonic development was observed to occur normally during the following hours post grafting (hpg), with labeled cells integrated in the embryo (**Figure 4C, D**). After fixation and methanol storage of chimeric embryos, FITC fluorescence in labelled grafted cells was completely lost (not shown), rendering necessary a revelation through immunohistochemistry. Fluorescent revelation of labeled cells with FITC-coupled tyramide was avoided because an extensive bleed through of fluorescence to channels at lower wavelengths was observed (not shown). Instead, revelation with TAMRA-coupled tyramide, whose fluorescent excitation/emission occurs at higher wavelengths (557nm and 583nm, respectively) than FITC (495nm and 521 nm, respectively), did not produce bleed through to lower wavelengths channels (**Figure 4G’, G” and H’, H”**).

**Figure 4.**
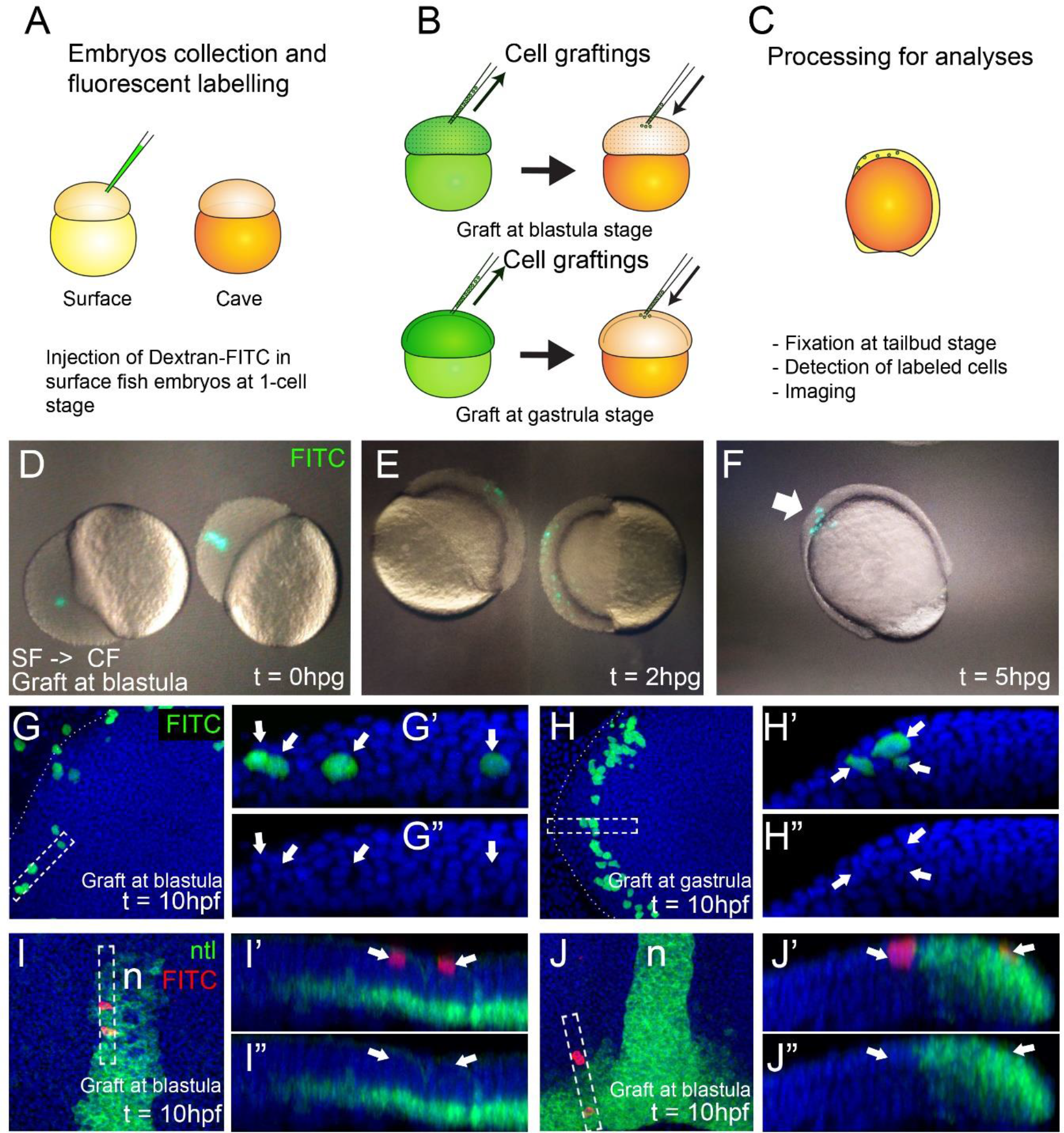
Generation of chimeric embryos in *A. mexicanus* by inter-morph cell transplantation. Procedure for cell grafting between surface and cavefish embryos (**A**). Live cavefish (host) embryos that have been transplanted with FITC-labeled surface cells (green) at the blastula stage, photographed immediately after the grafting (0 hpg, **B**), or after 2 hpg (**C**) and 5 hpg (**D**). Confocal images of dissected cavefish embryos at 10 hpf at the level of the anterior neural plate (indicated in dotted curved lines). Labeling in green corresponds to surface fish cells grafted at blastula (**E**) and gastrula (**F**) stages. Respective reconstructed sections were obtained at the levels indicated in rectangles (**E’, E”** and **F’, F”**). Confocal images of dissected cavefish embryos at 10 hpf stained for *ntl* (green), at the level of the anterior notochord (**G**) and the tail bud (**H**). Labeling in red corresponds to surface fish cells grafted at blastula stage. Arrows in **G’** and **G”** indicate superficial ectodermal cells overlying the notochord anterior tip (green). Arrowheads in J’ and J” indicate a cell in the mesodermal domain expressing *ntl*. Images in E, F G and H are dorsal views, anterior to the top. Corresponding reconstructions are oriented with anterior to the right and dorsal on top.

We compared the organization and repartition of grafted cells in host embryos at the two transplantation stages, blastula *versus* gastrula, and observed clear differences between the two conditions. Clones transplanted at blastula stage were distributed extensively throughout the embryos and in a disorganized and scattered manner. On the other hand, cells transplanted at gastrula stages were restricted in space and in some cases showed clear signs of symmetry (compare **Figure 4E** and **F**). Using the positional information and the expression of *ntl* as reference, we observed that cells grafted at blastula stages were able to produce both ectodermal (n = 15/15 embryos, superficial cells in **Figure 4G, G’, G”**) and endomesodermal (n = 6/15 embryos, *ntl* positive cell in **Figure 4H, H’, H”**) derivatives in chimeric embryos. On the other hand, grafts performed at gastrula stages gave rise to only ectodermal cells (n = 24/24 embryos, **Figure 4F** and not shown). These data were consistent with an expected progressive lineage restriction from mid-blastula to early gastrulation stages. Confocal reconstructions suggested a correct integration of transferred cells in the developing host tissues (**Figure 4E’, E”** and **F’, F”**). Similar results were also observed in reciprocal experiments, *i*.*e*. transplants of cavefish donor cells into a surface fish host (not shown). We also found that surface fish embryonic cells were able to differentiate into pigmented cells in a cavefish host (**Supplemental figure**) confirming the correct integration of donor cells in chimeric embryos and illustrating a typical cell-autonomous process.

Intermorph grafting will shed light on the cell autonomy and the effect of the embryonic signaling environment on previously described heterochronies, heterotopies and differences of gene expression levels of during development of *Astyanax* morphs (Hinaux et al., 2016; Pottin et al., 2011; Torres-Paz et al., 2019; Yamamoto et al., 2004). The combination of these grafting methods with the use of transgenic reporter lines such as the cavefish and surface fish *zic1:GFP* lines (Devos et al., 2019), will allow the detailed investigation of intrinsic and extrinsic factors implicated in eye specification and degeneration.

## Conclusions

Implementation and optimization of new methods in emergent model systems is fundamental for tackling novel scientific questions. Here we describe the methodology and potential applications of cellular techniques to generate yolk-free gastruloids and chimeric embryos in *Astyanax mexicanus*. These methods will allow the characterization of developmental states during cell lineages differentiation in embryogenesis. In addition, these techniques will push forward genomic and cellular approaches to understand the key steps during eye development and degeneration in cavefish.

## Acknowledgements

We thank Stéphane Père, Victor Simon and Krystel Saroul for taking care of our *Astyanax* colony, Maryline Blin for technical support for ISH protocols and François Agnès and all other current and past members of the DECA team for fruitful discussions and important suggestions. We thank also to Dr. Ben Steventon for his advices on the protocol for generation of pescoids, and Drs. Nicolas David and Lazaro Centanin for technical advices on cell transplantation methods. Grant support was received from UNADEV/AVIESAN/Vision and Retina France, CNRS and Becas Chile.

---

**Supplemental figure.**
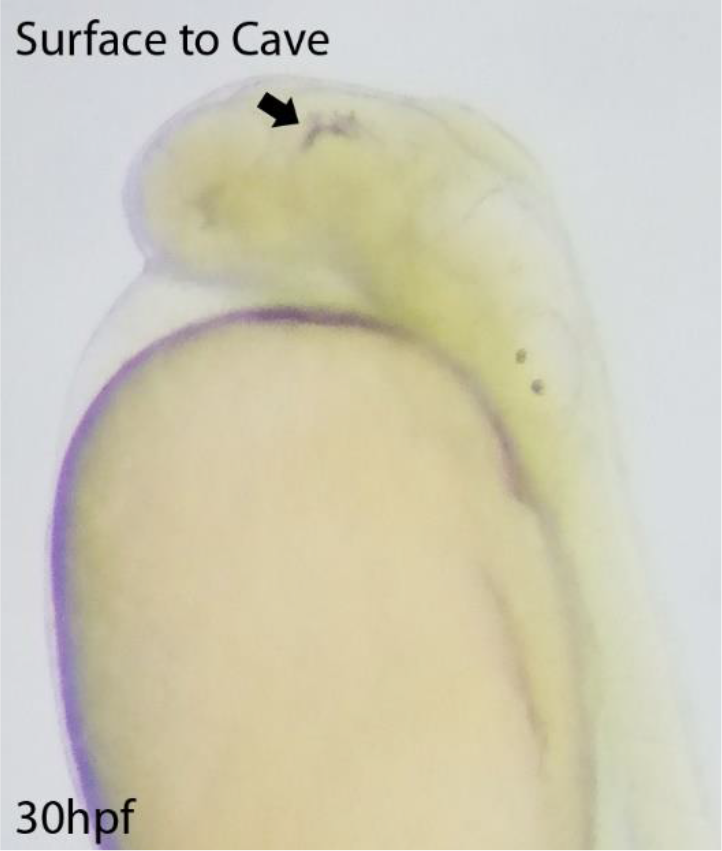
Differentiation of pigmented cells in chimeric embryo. Pigmented cell (arrow) derived from surface fish donor cells transplanted into a cavefish embryo at the blastula stage.

## Notes

### Competing Interest Statement

The authors have declared no competing interest.

## References

Alié, A., Devos, L., Torres-Paz, J., Prunier, L., Boulet, F., Blin, M., … Retaux, S. (2018). Developmental evolution of the forebrain in cavefish, from natural variations in neuropeptides to behavior. ELife, 7. https://doi.org/10.7554/eLife.32808

Bibliowicz, J., Alié, A., Espinasa, L., Yoshizawa, M., Blin, M., Hinaux, H., … Rétaux, S. (2013). Differences in chemosensory response between eyed and eyeless Astyanax mexicanus of the Rio Subterráneo cave, 2013(March), 2–7. https://doi.org/10.1186/2041-9139-4-25

Blin, M., Tine, E., Meister, L., Elipot, Y., Bibliowicz, J., Espinasa, L., & Rétaux, S. (2018). Developmental evolution and developmental plasticity of the olfactory epithelium and olfactory skills in Mexican cavefish. Developmental Biology, 441(2), 242–251. https://doi.org/10.1016/j.ydbio.2018.04.019

Cavodeassi, F., Carreira-Barbosa, F., Young, R. M., Concha, M. L., Allende, M. L., Houart, C., … Wilson, S. W. (2005). Early stages of zebrafish eye formation require the coordinated activity of Wnt11, Fz5, and the Wnt/β-catenin pathway. Neuron, 47(1), 43–56. https://doi.org/10.1016/j.neuron.2005.05.026

Cavodeassi, F., Ivanovitch, K., & Wilson, S. W. (2013). Eph/Ephrin signalling maintains eye field segregation from adjacent neural plate territories during forebrain morphogenesis. Development (Cambridge), 140(20), 4193–4202. https://doi.org/10.1242/dev.097048

Chan, S. H., Tang, Y., Miao, L., Darwich-Codore, H., Vejnar, C. E., Beaudoin, J. D., … Giraldez, A. J. (2019). Brd4 and P300 Confer Transcriptional Competency during Zygotic Genome Activation. Developmental Cell, 49(6), 867-881.e8. https://doi.org/10.1016/j.devcel.2019.05.037

Chen, S. R., & Kimelman, D. (2000). The role of the yolk syncytial layer in germ layer patterning in zebrafish. Development, 127(21), 4681–4689.

Devos, L., Klee, F., Edouard, J., Simon, V., Legendre, L., Khallouki, N. El, … Rétaux, S. (2019). Morphogenetic and patterning defects explain the coloboma phenotype of the eye in the Mexican cavefish. BioRxiv, 698035. https://doi.org/10.1101/698035

Elipot, Y., Legendre, L., Père, S., Sohm, F., & Rétaux, S. (2014). Astyanax Transgenesis and Husbandry: How Cavefish Enters the Laboratory. Zebrafish, 11(4), 291–299. https://doi.org/10.1089/zeb.2014.1005

Fauny, J.-D., Thisse, B., & Thisse, C. (2009). The entire zebrafish blastula-gastrula margin acts as an organizer dependent on the ratio of Nodal to BMP activity. Development, 136(22), 3811–3819. https://doi.org/10.1242/dev.039693

Fuhrmann, J. F., Buono, L., Adelmann, L., Martinez-Morales, J. R., & Centanin, L. (2020). Genetic developmental timing revealed by inter-species transplantations in fish. Development (Cambridge, England), 147(22). https://doi.org/10.1242/dev.192500

Fulton, T., Trivedi, V., Attardi, A., Anlas, K., Dingare, C., Arias, A. M., & Steventon, B. (2020). Axis Specification in Zebrafish Is Robust to Cell Mixing and Reveals a Regulation of Pattern Formation by Morphogenesis. Current Biology, 1–11. https://doi.org/10.1016/j.cub.2020.05.048

García-Calero, E., Fernández-Garre, P., Martínez, S., & Puelles, L. (2008). Early mammillary pouch specification in the course of prechordal ventralization of the forebrain tegmentum. Developmental Biology, 320(2), 366–377. https://doi.org/10.1016/j.ydbio.2008.05.545

Giger, F. A., Dumortier, J. G., & David, N. B. (2016). Analyzing in vivo cell migration using cell transplantations and time-lapse imaging in zebrafish embryos. Journal of Visualized Experiments, 2016(110), e53792. https://doi.org/10.3791/53792

Gordon, H. B., Lusk, S., Carney, K. R., Wirick, E. O., Murray, B. F., & Kwan, K. M. (2018). Hedgehog signaling regulates cell motility and optic fissure and stalk formation during vertebrate eye morphogenesis. Development (Cambridge), 145(22). https://doi.org/10.1242/dev.165068

Hinaux, H., Devos, L., Blin, M., Elipot, Y., Bibliowicz, J., Alié, A., & Rétaux, S. (2016). Sensory evolution in blind cavefish is driven by early embryonic events during gastrulation and neurulation. Development, 143(23), 4521–4532. https://doi.org/10.1242/dev.141291

Hinaux, H., Pottin, K., Chalhoub, H., Père, S., Elipot, Y., Legendre, L., & Rétaux, S. (2011). A Developmental Staging Table for Astyanax mexicanus Surface Fish and Pachón Cavefish. Zebrafish, 8(4), 155–165. https://doi.org/10.1089/zeb.2011.0713

Ivanovitch, K., Cavodeassi, F., & Wilson, S. W. (2013). Precocious acquisition of neuroepithelial character in the eye field underlies the onset of eye morphogenesis. Developmental Cell, 27(3), 293–305. https://doi.org/10.1016/j.devcel.2013.09.023

Jeffery, W. R. (2020). Astyanax surface and cave fish morphs. EvoDevo. BioMed Central. https://doi.org/10.1186/s13227-020-00159-6

Jesuthasan, S., & Strähle, U. (1997). Dynamic microtubules and specification of the zebrafish embryonic axis. Current Biology, 7(1), 31–42. https://doi.org/10.1016/S0960-9822(06)00025-X

Kimmel, C. B., Ballard, W. W., Kimmel, S. R., Ullmann, B., & Schilling, T. F. (1995). Stages of embryonic development of the zebrafish. Developmental Dynamics : An Official Publication of the American Association of Anatomists, 203(3), 253–310. https://doi.org/10.1002/aja.1002030302

Ma, L., Ng, M., Shi, J., Gore, A. V., Castranova, D., Weinstein, B. M., & Jeffery, W. R. (2020). Evolutionary Changes in Left-Right Visceral Asymmetry in Astyanax Cavefish. BioRxiv, 2020.05.15.098483. https://doi.org/10.1101/2020.05.15.098483

Ma, L., Strickler, A. G., Parkhurst, A., Yoshizawa, M., Shi, J., & Jeffery, W. R. (2018). Maternal genetic effects in Astyanax cavefish development. Developmental Biology, 441(2), 209–220. https://doi.org/10.1016/J.YDBIO.2018.07.014

Macdonald, R., Barth, K. a, Xu, Q., Holder, N., Mikkola, I., & Wilson, S. W. (1995). Midline signalling is required for Pax gene regulation and patterning of the eyes. Development (Cambridge, England), 121(10), 3267–3278. Retrieved from http://www.ncbi.nlm.nih.gov/pubmed/7588061

Marlow, F. L. (2020). Setting up for gastrulation in zebrafish. Current Topics in Developmental Biology (1st ed., Vol. 136). Elsevier Inc. https://doi.org/10.1016/bs.ctdb.2019.08.002

Pottin, K., Hinaux, H., & Rétaux, S. (2011). Restoring eye size in Astyanax mexicanus blind cavefish embryos through modulation of the Shh and Fgf8 forebrain organising centres, 2476, 2467–2476. https://doi.org/10.1242/dev.054106

Rétaux, S., Alié, A., Blin, M., Devos, L., Elipot, Y., & Hinaux, H. (2016). Neural Development and Evolution in Astyanax mexicanus: Comparing Cavefish and Surface Fish Brains. In Biology and Evolution of the Mexican Cavefish (pp. 227–244). Elsevier Inc. https://doi.org/10.1016/B978-0-12-802148-4.00012-8

Rodaway, A., Takeda, H., Koshida, S., Broadbent, J., Price, B., Smith, J. C., … Holder, N. (1999). Induction of the mesendoderm in the zebrafish germ ring by yolk cell-derived TGF-β family signals and discrimination of mesoderm and endoderm by FGF. Development, 126(14), 3067–3078.

Schauer, A., Pinheiro, D., Hauschild, R., & Heisenberg, C.-P. (2020). Zebrafish embryonic explants undergo genetically encoded self-assembly. ELife, 9. https://doi.org/10.7554/elife.55190

Schier, a F., Neuhauss, S. C., Helde, K. a, Talbot, W. S., & Driever, W. (1997). The one-eyed pinhead gene functions in mesoderm and endoderm formation in zebrafish and interacts with no tail. Development (Cambridge, England), 124(2), 327–342.

Schier, A. F., & Talbot, W. S. (2005). Molecular Genetics of Axis Formation in Zebrafish. Annual Review of Genetics, 39(1), 561–613. https://doi.org/10.1146/annurev.genet.37.110801.143752

Simon, V., Hyacinthe, C., & Rétaux, S. (2019). Breeding behavior in the blind Mexican cavefish and its river-dwelling conspecific. PLoS ONE, 14(2). https://doi.org/10.1371/journal.pone.0212591

Solnica-Krezel, L. (2020). Maternal contributions to gastrulation in zebrafish. Current Topics in Developmental Biology (1st ed., Vol. 140). Elsevier Inc. https://doi.org/10.1016/bs.ctdb.2020.05.001

Torres-Paz, J., Hyacinthe, C., Pierre, C., & Rétaux, S. (2018). Towards an integrated approach to understand Mexican cavefish evolution. Biology Letters, 14(8). https://doi.org/10.1098/rsbl.2018.0101

Torres-Paz, J., Leclercq, J., & Rétaux, S. (2019). Maternally-regulated gastrulation as a source of variation contributing to cavefish forebrain evolution. ELife, 8. https://doi.org/10.7554/eLife.50160

Varatharasan, N., Croll, R. P., & Franz-Odendaal, T. (2009). Taste bud development and patterning in sighted and blind morphs of Astyanax mexicanus. Developmental Dynamics, 238(12), 3056–3064. https://doi.org/10.1002/dvdy.22144

Vopalensky, P., Pralow, S., & Vastenhouw, N. L. (2018). Reduced expression of the nodal co-receptor Oep causes loss of mesendodermal competence in zebrafish. Development (Cambridge), 145(5). https://doi.org/10.1242/dev.158832

Williams, M. L. K., & Solnica-Krezel, L. (2020). Nodal and planar cell polarity signaling cooperate to regulate zebrafish convergence & extension gastrulation movements. ELife, 9, 1–25. https://doi.org/10.7554/eLife.54445

Woo, K., & Fraser, S. E. (1997). Specification of the zebrafish nervous system by nonaxial signals. Science, 277(5323), 254–257. https://doi.org/10.1126/science.277.5323.254

Yamamoto, Y, & Jeffery, W. R. (2000). Central role for the lens in cave fish eye degeneration. Science (New York, N.Y.), 289(5479), 631–633. https://doi.org/10.1126/SCIENCE.289.5479.631

Yamamoto, Yoshiyuki, Stock, D. W., & Jeffery, W. R. (2004). Hedgehog signalling controls eye degeneration in blind cavefish, 431(October). https://doi.org/10.1038/nature02906.1.

Yoshizawa, M., Gorički, Š., Soares, D., & Jeffery, W. R. (2010). Evolution of a behavioral shift mediated by superficial neuromasts helps cavefish find food in darkness. Current Biology, 20(18), 1631–1636. https://doi.org/10.1016/j.cub.2010.07.017

Yoshizawa, M., Hixon, E., & Jeffery, W. R. (2018). Neural Crest Transplantation Reveals Key Roles in the Evolution of Cavefish Development. Integrative and Comparative Biology, 58(3), 411–420. https://doi.org/10.1093/icb/icy006

Zuber, M. E., Gestri, G., Viczian, A. S., Barsacchi, G., & Harris, W. A. (2003). Specification of the vertebrate eye by a network of eye field transcription factors, 5155–5167. https://doi.org/10.1242/dev.00723

